# The retromer complex regulates *C. elegans* development and mammalian ciliogenesis

**DOI:** 10.1101/2021.09.24.461716

**Authors:** Shuwei Xie, Ellie Smith, Carter Dierlam, Danita Mathew, Angelina Davis, Allana Williams, Naava Naslavsky, Jyoti Iyer, Steve Caplan

**Author notes:** To whom correspondence should be addressed: Jyoti Iyer: ^2^Department of Chemistry and Biochemistry, University of Tulsa, Tulsa, OK 74104., Steve Caplan: ^1^Department of Biochemistry & Molecular Biology, University of Nebraska Medical Center, Omaha NE 68198.

## Abstract

The mammalian retromer is comprised of subunits VPS26, VPS29 and VPS35, and a more loosely-associated sorting nexin (SNX) heterodimer. Despite known roles for the retromer in multiple trafficking events in yeast and mammalian cells, its role in development is poorly understood, and its potential function in primary ciliogenesis remains unknown. Using CRISPR-Cas9 editing, we demonstrated that *vps-26* homozygous knockout *C. elegans* have reduced brood sizes and impaired vulval development, as well as decreased body length which has been linked to defects in primary ciliogenesis. Since many endocytic proteins are implicated in the generation of primary cilia, we addressed whether the retromer regulates ciliogenesis in mammalian cells. We observed VPS35 localized to the primary cilium, and depletion of VPS26, VPS35 or SNX1/SNX5 led to decreased ciliogenesis. Retromer also coimmunoprecipitated with the capping protein, CP110, and was required for its removal from the mother centriole. Herein, we characterize new roles for the retromer in *C. elegans* development and in the regulation of ciliogenesis in mammalian cells, and suggest a novel role for the retromer in CP110 removal from the mother centriole.

## Introduction

The retromer is a conserved protein complex involved in the sorting and trafficking of endosomal cargo (Naslavsky and Caplan, 2018; Wang et al., 2018). Studies describe the mammalian retromer as being comprised of a “core complex” of the VPS26/VPS29/VPS35 trimer (Haft et al., 2000). This core complex binds to a heterodimeric sorting nexin duo comprised of either SNX1/SNX2 (Griffin et al., 2005; Rojas et al., 2007) or SNX5/SNX6 (Cullen and Korswagen, 2011; Haft et al., 2000). However, evidence in mammals suggests that the SNX protein association with the retromer core is less stable than in yeast (Harbour et al., 2010) and that the SNX dimers can function independently of the core complex (Kvainickas et al., 2017). While the sorting nexin proteins bind to endosomal-enriched phosphatidylinositol-3-phosphate via their phox domains (Seaman and Williams, 2002), the original function identified for the retromer core complex proteins was in binding to select cargo receptors and the regulation of their sorting/trafficking (Arighi et al., 2004; Seaman, 2004). Newer studies support the notion that sorting nexins, including the more peripherally associated SNX27, SNX17 and SNX3 proteins, also interact with cargo receptors and play an important role in the cargo selection process (Clairfeuille et al., 2016; Farfan et al., 2013; Seaman, 2021; Steinberg et al., 2012; van Kerkhof et al., 2005). The retromer was originally described for its role in endosomal retrieval of the carboxypeptidase Y (CPY) hydrolase receptor, Vps10p, back to the Golgi after it delivers CPY to the yeast vacuole (Seaman et al., 1997), as well as its function in the retrieval of the cation-independent mannose-6-phosphate receptor from endosomes to the Golgi in mammalian cells (Arighi et al., 2004). However, recent studies indicate that the retromer plays a central role in many additional cellular trafficking events (Seaman, 2021; Wang et al., 2018).

The retromer core complex of VPS26/VPS29/VPS35, whose structure has recently been elucidated (Chen et al., 2019; Norwood et al., 2011), is highly conserved throughout the evolutionary ladder from yeast, to invertebrates such as *C. elegans*, as well as humans (Wang et al., 2018). For example, there is 58% identity shared between human VPS26A (one of the two VPS26 isoforms evolved from duplicated genes (Bugarcic et al., 2011; Kerr et al., 2005)) and the *C. elegans* VPS-26 proteins, 58% identity shared between human VPS29 and the *C. elegans* VPS-29 proteins, and 49% identity between the human VPS35 and the worm VPS-35 proteins. Whereas the role of the retromer complex has been studied extensively in yeast and in mammalian systems, the function of retromer and its role in development in invertebrates such as *C. elegans* is incompletely understood. It has been observed that worms with mutant *vps-29* or *vps-35* genes display impaired trafficking of the receptor-type guanylate cyclase GCY-9, which accumulates in BAG chemosensory neurons (Coudreuse et al., 2006; Martinez-Velazquez and Ringstad, 2018). Recent studies have shown a role for worm retromer subunits in anteroposterior polarity, Wnt signaling and Q cell migration (Prasad and Clark, 2006). Furthermore, *vps-35* knockout worms have reduced mTorc1 signaling and increased lifespans (Kvainickas et al., 2019). Nonetheless, important questions remain with regard to the role of retromer in the *C. elegans* nematode model system, particularly with regard to development.

The retromer complex has been documented as having a wide and growing array of functions in recent years. In addition to membrane trafficking and the regulation of mannose-6-phosphate receptor recovery from endosomes to the Golgi (Arighi et al., 2004; Reddy and Seaman, 2001; Seaman, 2004), and its role in receptor recycling, endosomal tubule dynamics and modulation of the actin cytoskeleton through FAM21 and the WASH complex (Gokool et al., 2007; Gomez and Billadeau, 2009; Harbour et al., 2010; Strochlic et al., 2007), the retromer has also been implicated in apoptosis (Farmer et al., 2019), mitochondrial membrane dynamics and Parkinson’s disease (Braschi et al., 2010; Farmer et al., 2018; Farmer et al., 2017; Kumar et al., 2012; Vilarino-Guell et al., 2011; Wang et al., 2016; Wang et al., 2017; Zimprich et al., 2011), as well as in the regulation of centrosome duplication (Xie et al., 2018). Despite the high level of retromer conservation across species, surprisingly, the role of retromer and its function in the model organism *C. elegans* has not been studied extensively. Moreover, despite the interaction of retromer with EHD proteins and their binding partner (Gokool et al., 2007; McKenzie et al., 2012; Zhang et al., 2012a), MICAL-L1 (Zhang et al., 2012b), and the known involvement of the latter proteins in regulating primary ciliogenesis (Lu et al., 2015; Xie et al., 2019), to date the involvement of retromer proteins in primary cilia biogenesis remains to be elucidated.

Herein, we used CRISPR-based technology to first obtain a *vps-26::ha* tagged *C. elegans* strain to detect endogenous VPS-26 expression in worms. Using this strain as a background, we again used CRISPR to successfully knockout VPS-26 expression. We demonstrate that while *vps-26* homozygous knockout worms are no less viable than their wild-type counterparts, the knockout worms have dramatically reduced brood sizes typically with 10-fold fewer progeny. Impaired vulval development was observed in the *vps-26* knockout worms which could explain why these worms produce fewer progeny. In addition to decreased brood sizes, the *vps-26* knockout adult worms also display a significant decrease in body length, a characteristic previously linked to defects in primary ciliogenesis. Given the known involvement of retromer interaction partners such as EHD proteins and MICAL-L1 in regulating ciliogenesis, we assessed the role of retromer subunits in primary ciliogenesis in RPE-1 cells. SiRNA-based knockdown of VPS26, VPS35 or the SNX1 and SNX5 sorting nexins led to reduced levels of primary ciliogenesis. Despite retromer interaction with EHD proteins and MICAL-L1, the retromer subunits were not required for the localization of either MICAL-L1 or EHD1 to primary cilia. Conversely, the retromer subunit VPS35 was recruited to primary cilia independently of MICAL-L1 and EHD1. Finally, we demonstrate that depletion of either VPS26 or VPS35 leads to the failure of CP110 removal from the mother centriole, which is a key step in primary ciliogenesis. Moreover, we provide initial evidence of a complex formed between CP110 and the retromer subunits, hinting at a potential explanation for the elusive mechanism of CP110 removal from the mother centriole as ciliogenesis progresses. Overall, our study highlights new roles for the retromer complex in *C. elegans* development and in the regulation of primary ciliogenesis in mammalian cells, expanding on the broad function of retromer and providing new clues to the mechanism of CP110 removal from the mother centriole.

## RESULTS

### Generation of CRISP/Cas9 gene-edited *vps-26* worm strains

To investigate the role of the retromer complex in *C. elegans*, we probed the function of the conserved retromer complex subunit VPS-26 which shares 58% identity with the human VPS26A subunit. Indeed, it has been demonstrated that knockdown of VPS26 expression in mammalian cells leads to destabilization of VPS35, thus disrupting the entire retromer complex (Seaman, 2004). The lack of an available antibody to the *C. elegans* VPS-26 protein led us to generate a worm strain expressing epitope-tagged VPS-26. To study the expression and localization of *C. elegans* VPS-26, CRISPR/Cas9 genome editing was performed to create a strain expressing VPS-26 fused to a C-terminal hemagglutinin (HA) tag. The schematic of the repair template and the generated *vps-26::ha* strain is shown in Fig 1A. To facilitate screening of positively edited worms, a NdeI restriction site was introduced into the repair template (Fig. 1A). Positive F1 heterozygotes were first identified by PCR and restriction digestion followed by agarose gel electrophoresis. Subsequently, we again screened the progeny of the positively edited heterozygous worms for homozygosity of the edit (Fig. 1B). As shown in Fig. 1B, homozygous-edited *vps-26::ha* worms could be successfully identified. The identified homozygotes were further confirmed by DNA sequencing (Fig. 1C).

**Fig. 1.**
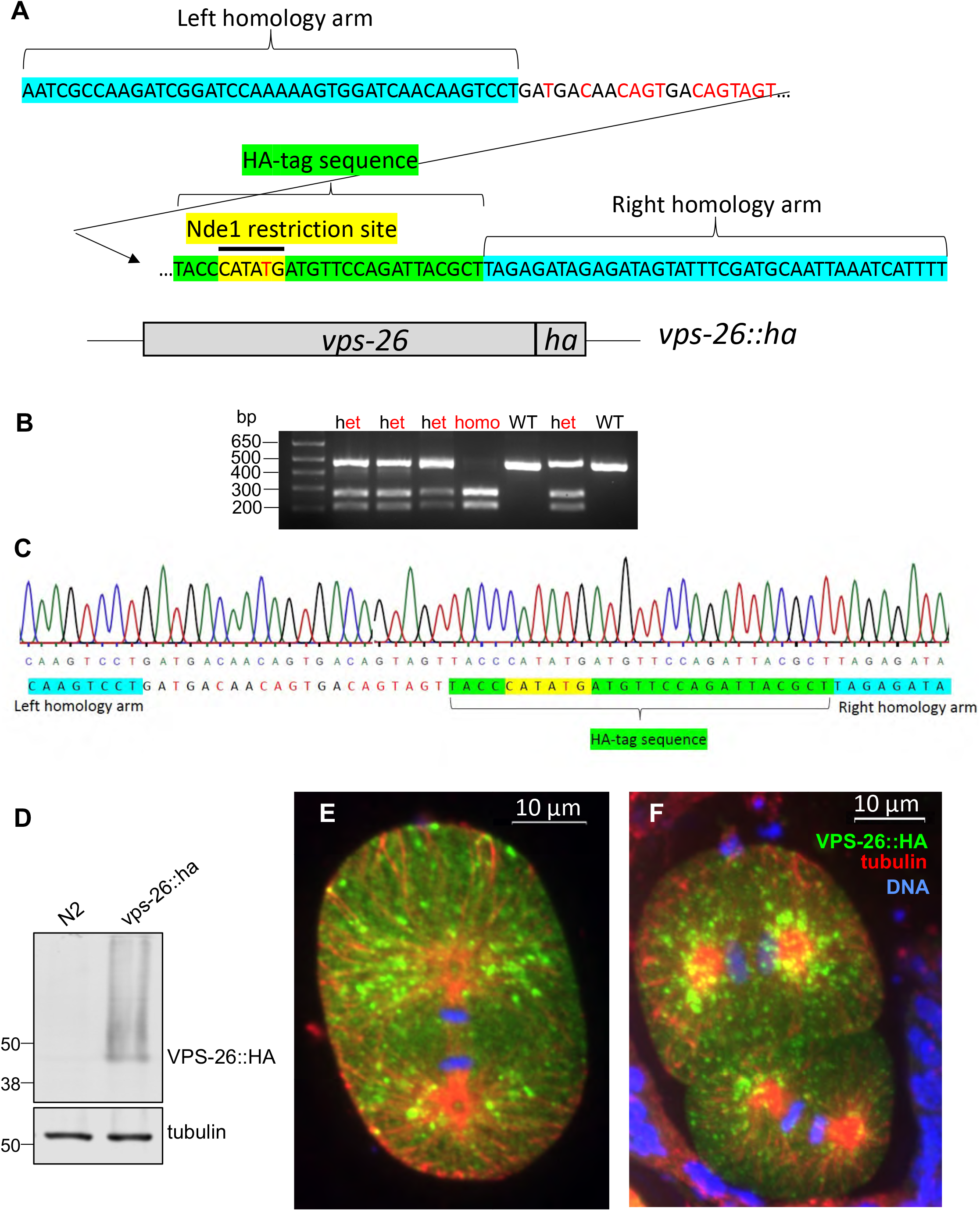
Expression and validation of VPS-26 expression in *C. elegans*. (A) Schematic of the CRISPR repair template used for inserting a (hemagglutinin-tag) HA-tag sequence at the 3’ end of the *C. elegans vps-26* gene. (B) Agarose gel electrophoresis of NdeI digested PCR products to screen for homozygous-edited *C. elegans* carrying the *vps-26::ha* edit. het: heterozygote; homo: homozygote; WT: wild-type. (C) Alignment of sequencing data from an identified *vps-26::ha* homozygous worm line with the designed CRISPR repair template demonstrating successful introduction of the HA-tag DNA sequence at the 3’end of the *vps-26* gene. (D) Immunoblot analysis showing VPS-26::HA expression in homozygous edited worms (right lane). The wild-type N2 strain served as a negative control for the experiment (left lane). Alpha tubulin was used as the loading control. (E) and (F) Immunofluorescence staining showing VPS-26::HA expression in early *C. elegans* embryos. Green: VPS-26::HA; Red: Microtubules stained with alpha tubulin; Blue: DNA (E) VPS-26::HA expression in a 1-cell anaphase *C. elegans* embryo (F) VPS-26::HA expression in a 2-cell *C. elegans* embryo. The anterior cell is in telophase while the posterior cell is in anaphase. Scale bars, 10 μm.

Once the *vps-26::ha* homozygotes were confirmed, immunoblotting was performed and we detected abundant VPS-26::HA protein expression in *C. elegans* whole worm lysates (Fig. 1D). We then immunostained early embryos expressing VPS-26::HA to characterize the localization pattern of endogenously expressed VPS-26 (Fig. 1E and 1F). Accordingly, immunofluorescence analysis demonstrated that VPS-26::HA exhibits a punctate localization pattern around the spindle microtubules in 1-cell and 2-cell stage *C. elegans* embryos in mitosis (Fig. 1E and 1F).

To study the function of the retromer complex in *C. elegans*, CRISPR/Cas9 genome editing was used to knockout VPS-26 protein expression (Fig. 2). Specifically, two premature stop codons separated by a single base were introduced into the CRISPR repair template to cause a frameshift mutation (Fig. 2A). The schematic for the CRISPR repair template used to knockout VPS-26 expression is shown in Figure 2A. The *vps-26::ha* strain characterized in Fig. 1 was used as the background strain to knockout VPS-26 expression. Upon identifying homozygous-edited worms, DNA sequencing was performed to confirm successful integration of the CRISPR edit into the *vps-26* gene (Fig. 2B and 2C). Alignment of the sequencing data with the wild-type *vps-26* gene sequence demonstrated that a part of the *vps-26* gene sequence was deleted and two premature stop codons and the frameshift mutation were successfully inserted at the 5’ end of the *vps-26* gene sequence (Fig. 2B). Immunoblot analysis of whole worm lysates showed that VPS-26::HA expression was undetectable in the *vps-26* knockout strain as compared with the control *vps-26::ha* strain (Fig. 2D). This *vps-26* knockout strain was named *vps-26(luv21*). For the sake of simplicity, the *vps-26(luv21*) strain has been referred to as the *vps-26* knockout strain throughout this manuscript.

**Fig. 2.**
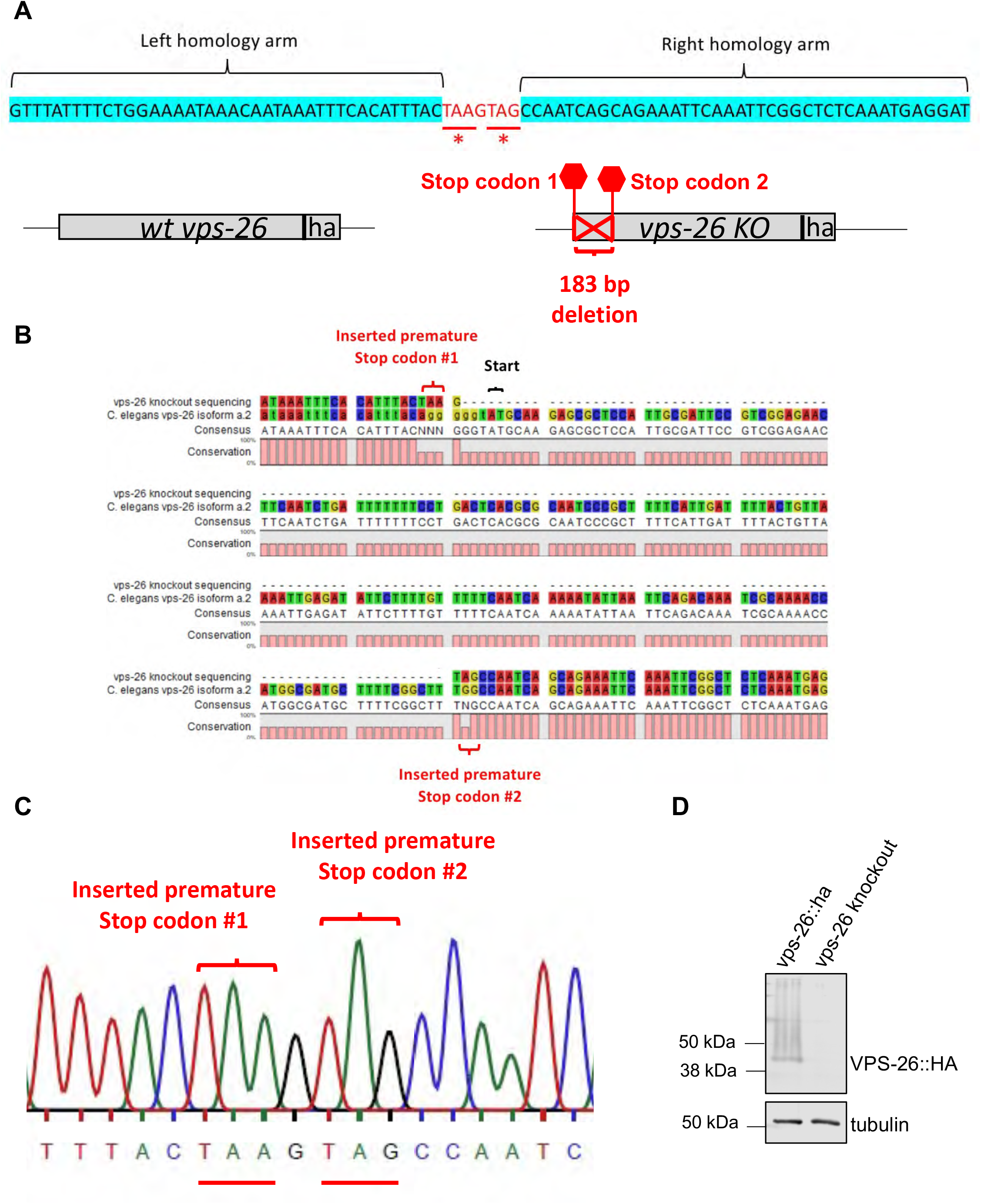
Deletion of *vps-26* in *C. elegans*. (A) Schematic of the CRISPR repair template used to knockout *C. elegans* VPS-26. (B) Alignment of sequencing data obtained from an identified *vps-26* deletion homozygote with the wild-type *vps-26* gene sequence confirming the insertion of the designed repair template into the *vps-26* gene. (C) Sequencing data obtained from *vps-26* knockout worms demonstrating successful insertion of the two premature stop codons into the *vps-26* gene via the CRISPR repair template. (D) Western blot showing that VPS-26::HA protein expression was undetectable in the *vps-26* knockout worms indicating that the knockout was successfully generated. The *vps-26::ha* strain served as a positive control for the experiment and alpha tubulin was used as the loading control.

### *vps-26* knockout worms have a decreased number of progeny, but normal viability

To determine the effect of depleting the retromer complex subunit VPS-26 on *C. elegans* development, brood counts and embryonic viability assays were done. Brood count assays performed at 20°C showed that *vps-26* knockout worms had a markedly reduced mean brood size of 18 (n=50) as compared with a mean brood size of 207 in control *vps-26::ha* worms (n=20) (2-tailed unpaired t-test, p<0.0001) (Fig. 3A).

**Fig. 3.**
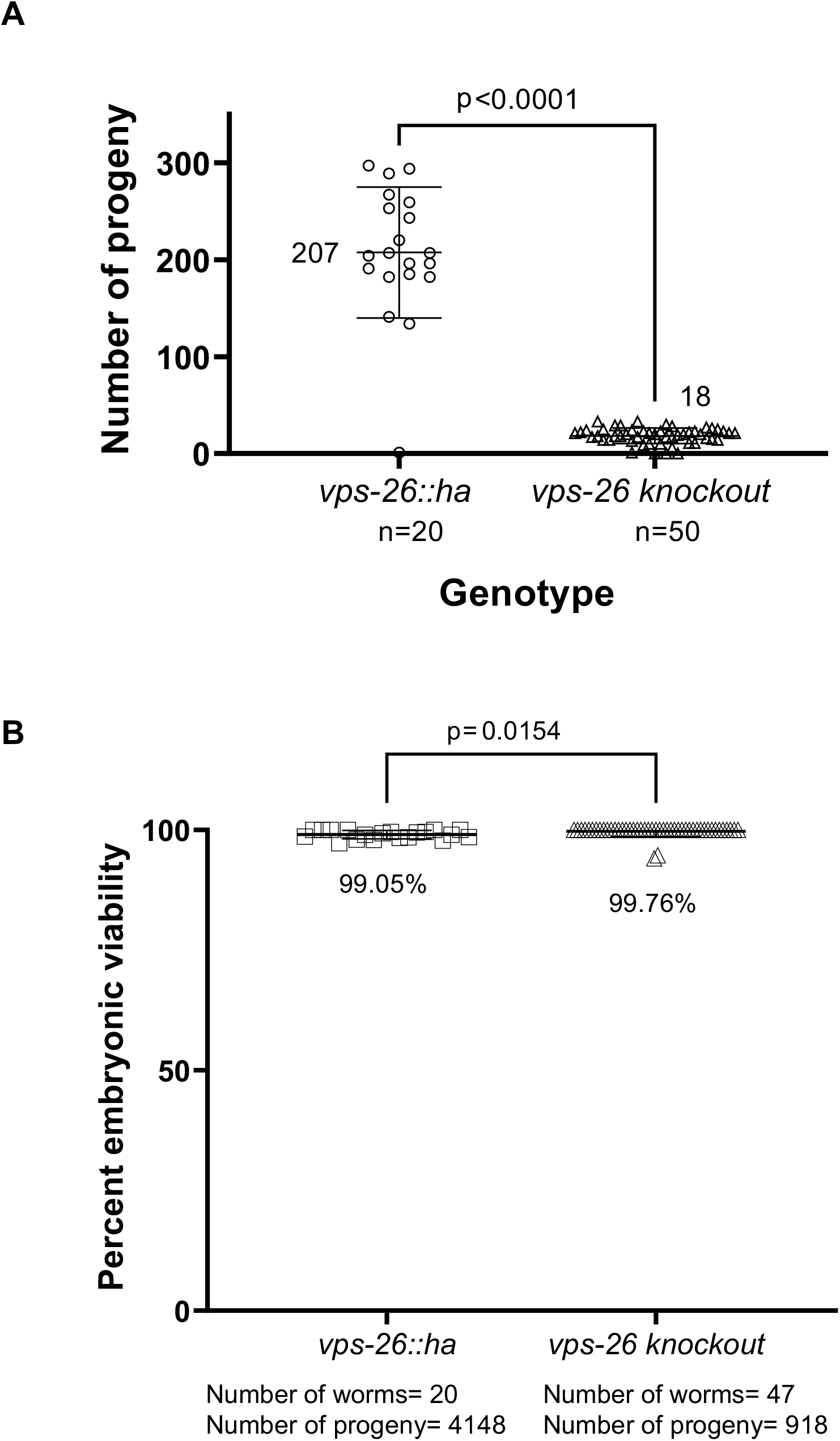
Effect of *vps-26* knockout on brood size and embryonic viability. (A) Quantification of brood sizes of *vps-26::ha* and *vps-26* knockout worms. The *vps-26* knockout worms exhibited a significantly reduced average brood size of 18 compared to an average brood size of 207 by control *vps-26::ha* worms at 20°C. p<0.0001 (unpaired t-test). n= Number of worms whose brood was analyzed for each genotype. (B) Quantification of embryonic viability of *vps-26::ha* and *vps-26* knockout worms. No significant change in embryonic viability was observed at 20°C upon the depletion of VPS-26. p=0.0154 (unpaired t-test). Error bars represent the standard deviation.

Although the brood size of *vps-26* knockout worms was reduced by more than 10-fold compared to the control strain, nevertheless almost all the eggs that were laid by the *vps-26* knockout worms were viable (Fig. 3B). Specifically, the mean embryonic viability of *vps-26* knockout worms was 99.76% (number of progeny analyzed=918) which was almost indistinguishable from the mean embryonic viability of the control *vps-26:ha* worms which was 99.05% (number of progeny analyzed=4148) (2-tailed unpaired t-test, p=0.0154) (Fig. 3B). These data indicate that VPS-26 function is critical for the maintenance of normal brood size but dispensable for embryonic viability.

### *vps-26* knockout worms display decreased body length compared to normal counterparts

We next examined whether defective vulva development might be responsible for the significantly reduced brood size observed in the *vps-26* knockout worms. Upon differential interference contrast imaging of control and *vps-26* knockout worms, we observed that the knockout worms frequently displayed defects in vulva development (Fig. S1). Specifically, a high proportion of the *vps-26* knockout worms displayed a protruding vulva which precluded them from laying eggs (Fig. S1B). In worms with a protruding vulva, this likely leads to the build-up and subsequent hatching of eggs inside their uterus, which subsequently causes the parent worm to die, thereby reducing the overall brood size.

Initial cursory examination of *vps-26* knockout worms through a dissecting microscope suggested that they may be smaller in size than their wild-type counterparts. Therefore, a thorough characterization of wild-type and *vps-26* knockout worm lengths was performed. The mean body length of the *vps-26* knockout worms was 475 µm (n=37) compared to 524 µm (n=30) for the control *vps-26::ha* worms (n=30) (2-tailed unpaired t-test, p<0.0001). These data indicate that VPS-26 function is essential for development and/or maintenence of normal body length in *C. elegans*.

A previous study indicated that decreased body size in *C. elegans* is often associated with ciliogenesis defects (Fujiwara et al, 2002). Moreover, since the retromer binding partners EHD1 (Gokool et al., 2007) and MICAL-L1 (Zhang et al., 2012a; Zhang et al., 2012b) have already been implicated in regulating ciliogenesis in mammalian cells (Lu et al, 2015; Xie et al, 2019), we then questioned whether the retromer complex is required for mammalian ciliogenesis.

### Retromer complex subunits are required for normal primary ciliogenesis

To determine if the retromer complex is required for the generation of the primary cilium, we first depleted RPE-1 cells of two different retromer core complex subunits, either VPS26 or VPS35, using siRNA oligonucleotides (Fig. 5D). We then induced primary ciliogenesis by serum-starvation and assessed the generation of the cilia after marking them with an antibody against acetylated-tubulin (Fig. 5A-C). While ∼50% of Mock-treated RPE-1 cells formed primary cilia (Fig. 5A, see arrows marking cilia, and quantified in Fig. 5E), reduced expression of either VPS26 or VPS35 resulted in a greater than two-fold decrease in the number of ciliated RPE-1 cells (Fig. 1B and Fig. 1C, quantified in E).

**Fig. 4.**
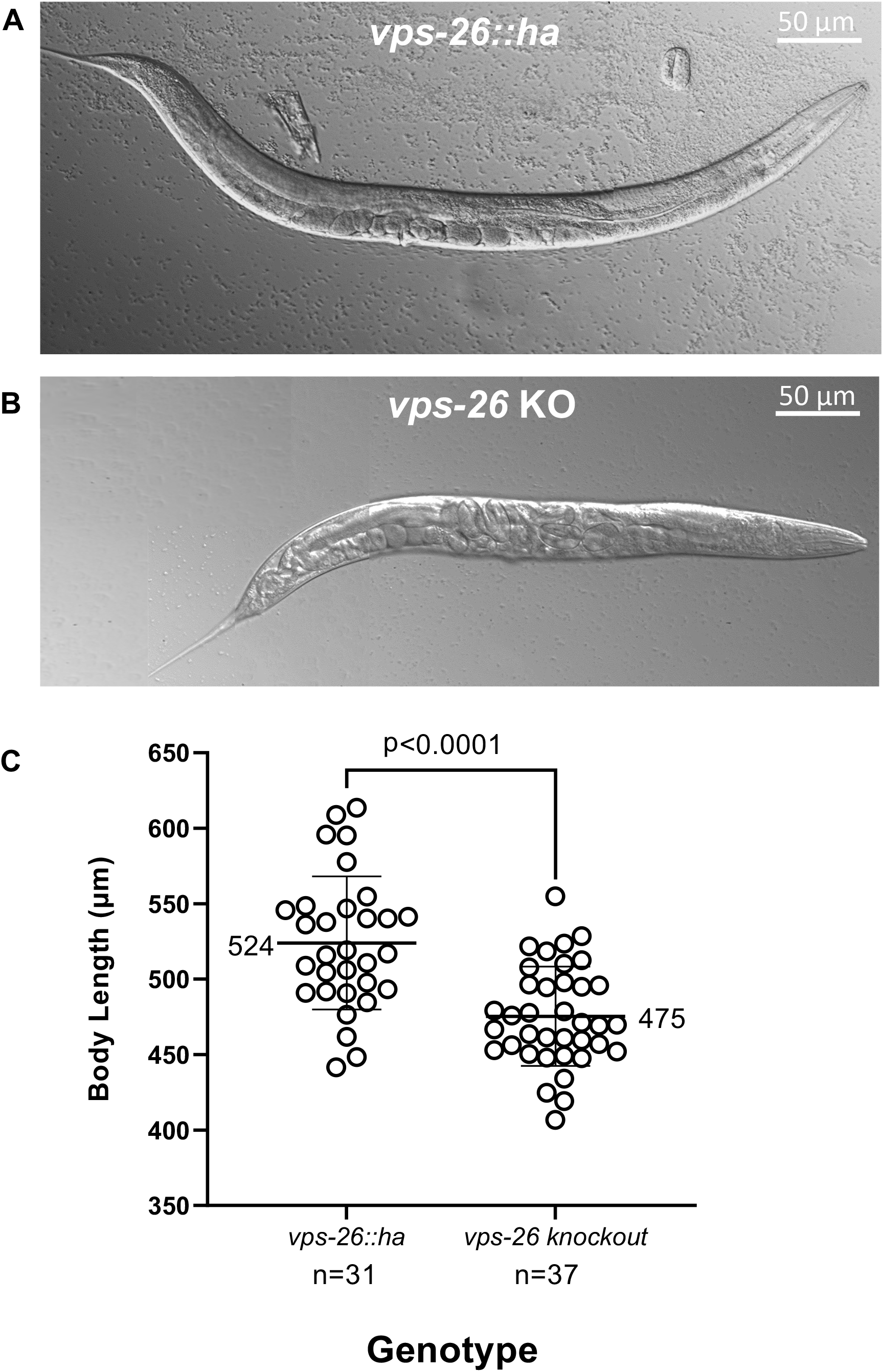
Effect of *vps-26* knockout on *C. elegans* body length. (A) A representative differential interference contrast (DIC) image showing the body length of a control VPS-26::HA worm. Scale bar: 50 µm (B) A representative DIC image showing the body length of a *vps-26* knockout worm. Scale bar: 50 µm (C) Quantification of body lengths of *vps-26::ha* and *vps-26* knockout worms. *vps-26* knockout worms have a shorter body length than *vps-26::ha* worms. The mean body length of *vps-26* knockout worms is 475 µm compared to 524 µm for control *vps-26::ha* worms. p<0.0001 (unpaired t-test). n= Number of independent worms whose body lengths were measured for each genotype. Error bars represent the standard deviation.

**Fig. 5.**
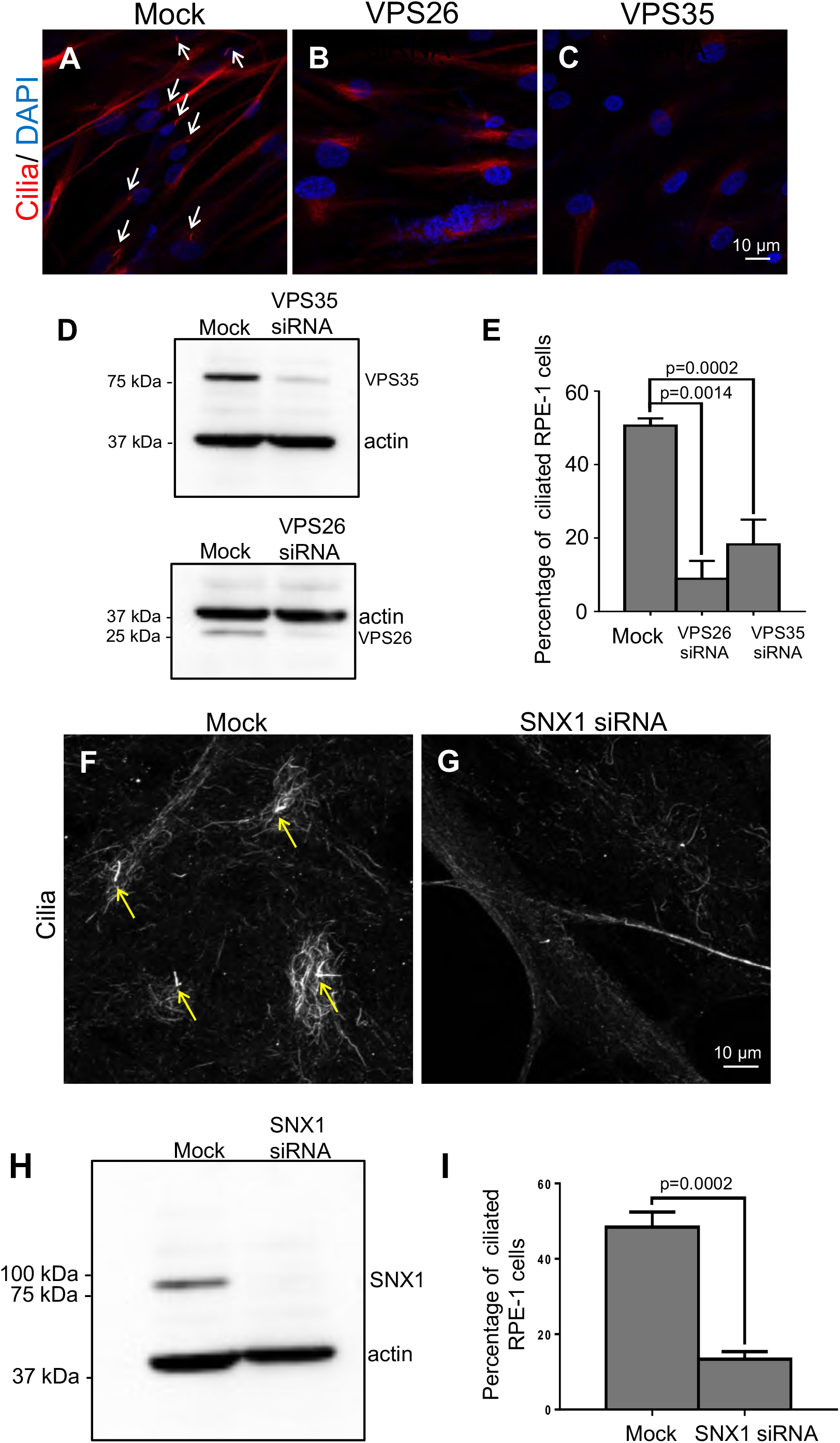
VPS26, VPS35 and SNX1 are required for normal ciliogenesis. (A-E) RPE-1 cells were Mock-treated (with lipofectamine RNAi-MAX in the absence of siRNA oligonucleotides), or treated with VPS26 or VPS35 siRNA for 48 h. Cells were then serum-starved for 24 h to induce ciliogenesis. Immunofluorescence staining of acetylated-tubulin reveals cilia (marked by white arrows). Compared to Mock-treated cells (A), fewer cilia were generated upon depletion of VPS26 (B) or VPS35 (C). VPS26 and VPS35 siRNA-depletion efficacies were determined by immunoblotting (D). The percentages of ciliated RPE-1 cells from either Mock- or siRNA-treated cover-slips were quantified and presented as a bar graph (E). (F-I) RPE-1 cells were either Mock-treated or treated with SNX1-siRNA for 48 h, followed by serum-starvation for 24 h to induce ciliogenesis. Cells were then fixed and immunostained for acetylated-tubulin to reveal cilia (denoted by yellow arrows). Compared to Mock (F), fewer cilia were detected in cells depleted of SNX1 (G). SNX1 knockdown efficacy was demonstrated by immunoblotting (H), and the percentage of ciliated Mock- or SNX1-siRNA depleted cells was quantified and presented as a bar graph (I). p-values were calculated for comparison between Mock- and siRNA-treated cells; *n*=3 experiments (>100 cells quantified for each experiment). Error bars denote standard deviation.

Since in mammalian cells the VPS26/VPS29/VPS35 heterotrimer loosely binds to the sorting nexin dimer, which can function independently of the core complex (Kvainickas et al., 2017; Simonetti et al., 2017), we also examined the role of the affiliated SNX1 protein in ciliogenesis using siRNA (Fig. 5F-I). Indeed, whereas ∼50% of Mock-treated cells were ciliated (Fig. 5F, see arrows marking cilia; quantified in Fig. 5I), upon SNX1-depletion only ∼10% of cells were detected with cilia (Fig. 5G; quantified in Fig. 5I). These data provide evidence that the retromer complex, including the VPS26/VPS29/VPS35 heterotrimer and affiliated SNX1 dimer, is required for ciliogenesis in mammalian cells.

### The retromer complex localizes to the primary cilium

Given the involvement of the retromer in ciliogenesis, we next assessed whether the retromer localizes to the membrane of the primary cilium and/or ciliary pocket. To address this, serum-starved RPE-1 cells were immunostained with antibodies to acetylated-tubulin to mark cilia, and with antibodies to VPS35 to evaluate the localization of the retromer complex (Fig. 6A). In ciliated cells, endogenous VPS35 was indeed detected at the base of the cilium (arrowhead), potentially at the ciliary pocket membrane, as well as on the daughter centriole (arrow), but not along the length of the cilium (Fig. 6A, dashed oval marks the centrioles).

**Fig. 6.**
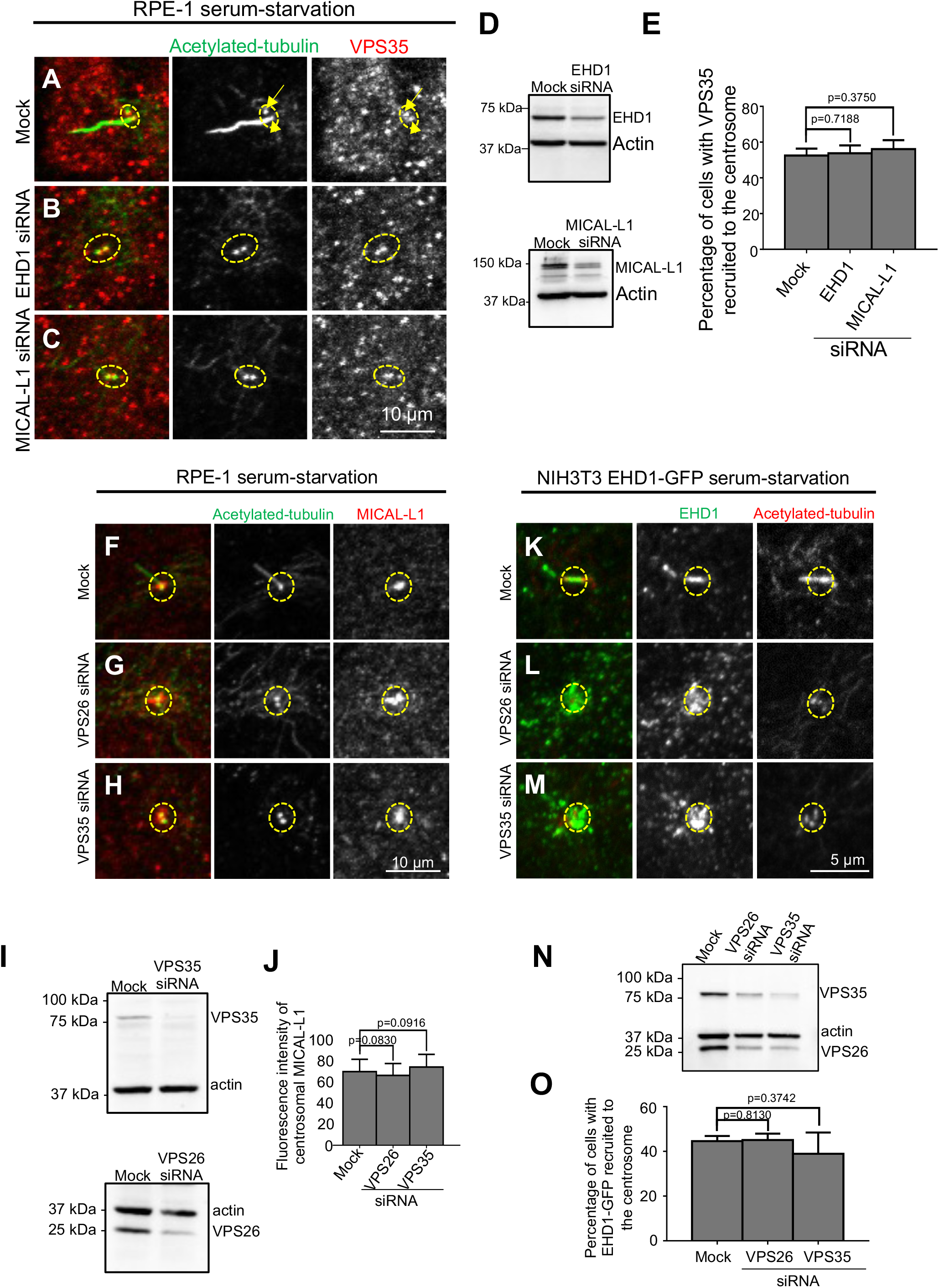
VPS35 is recruited to the basal body/centrioles independently of MICAL-L1 and EHD1. (A-E) RPE-1 cells were Mock-treated (A), EHD1-siRNA-treated (B), or MICAL-L1-treated (C), and then serum-starved for 1 h to induce ciliogenesis. Cells were fixed and immunostained with antibodies to acetylated-tubulin to mark cilia and/or centrioles (green) and VPS35 (red). VPS35 localized to both mother (yellow arrowhead) and daughter centrioles (yellow arrow) in Mock-treated and siRNA-treated cells (A-C, dashed ovals). The efficacy of EHD1 and MICAL-L1 knock-down was determined by immunoblot (D). Bar graph representing the percentage of cells with VPS35 detectable at the centrosome in Mock-treated cells, or cells depleted of EHD1 or MICAL-L1 (E). **Retromer is non-essential for the recruitment MICAL-L1 and EHD1 to the centrosome**. (F-J) Mock-treated (F), VPS26-siRNA-treated (G), or VPS35-siRNA-treated (H) RPE-1 cells were serum-starved for 1 h to induce ciliogenesis. The cells were then fixed and immunostained for acetylated-tubulin (green) to reveal cilia and/or centrioles, and for MICAL-L1 (red) to reveal its localization (F-H). MICAL-L1 was detected on acetylated-tubulin-labeled cilia/centrioles in both Mock-treated (F) and VPS26-(G) or VPS35-depleted (H) cells. VPS35- and VPS26-depletion by siRNA was demonstrated by immunoblotting (I). A circular region of interest (ROI) of 1.57 µm in diameter was projected around the basal body or mother centriole. The fluorescence intensity of the MICAL-L1 immunostaining within this ROI was measured and is presented as a bar graph (J). (K-O) CRISPR/Cas9 gene-edited NIH3T3 cells expressing EHD1-GFP were Mock-treated (K) or either VPS26-siRNA-treated (L), or VPS35-siRNA-treated (M), serum-starved for 1 h, and immunostained for acetylated-tubulin (red) or with anti-GFP antibodies to reveal EHD1 localization (green). EHD1 localized to acetylated-tubulin-labeled cilia/centrioles in Mock-treated (K) and VPS26-depleted (L) or VPS35-depleted (M) cells. VPS35- and VPS26-depletion in NIH3T3 gene-edited cells by siRNA was demonstrated by immunoblotting (N). Recruitment of EHD1 to the centrosome was determined by projecting a ROI of 1.57 µm in diameter around the basal body or mother centriole and measuring the fluorescence intensity of EHD1 immunostaining within this ROI (O). *n*=3 experiments (>100 cells quantified for each experiment). Yellow dshaed ovals or circles denote the ROIs and region of the centrioles. Error bars denote standard deviation.

### Recruitment of VPS35 to the centrosome is independent of MICAL-L1 and EHD1

Both EHD1 and MICAL-L1 interact with the retromer complex (Gokool et al., 2007; Zhang et al., 2012a; Zhang et al., 2012b). MICAL-L1 is recruited to the centrosome through an interaction with tubulin, and EHD1 is recruited by MICAL-L1 (Xie et al., 2019). Both MICAL-L1 and EHD1 are required for the generation of primary cilia (Lu et al., 2015; Xie et al., 2019). Since we observed localization of VPS35 to the primary cilium, we asked whether the retromer is recruited onto the ciliary base and/or the centrioles through its association with MICAL-L1 and/or EHD1. Accordingly, we used siRNA to deplete RPE-1 cells of MICAL-L1 or EHD1 (Fig. 6D), and immunostained them for acetylated-tubulin and endogenous VPS35 (Fig. 6B and C). As expected upon serum-starvation and previously published, ciliogenesis was impaired in the absence of EHD1 or MICAL-L1 (Lu et al., 2015; Xie et al., 2019) (compare Fig. 6A to Fig. 6B and Fig. 6C). However, despite the lack of a primary cilium in most cells lacking EHD1 or MICAL-L1, VPS35 was nonetheless observed at the centrioles of these non-ciliated cells (Fig. 6B,C and E; dashed ovals mark the VPS35 localized to centrioles).

Given the consistent localization of VPS35 to the mother and daughter centrioles, we questioned whether the retromer might be required for recruitment of MICAL-L1 and EHD1 to the centrioles and primary cilium. Accordingly, we used siRNA oligonucleotides to deplete expression of either VPS35 or VPS26 in both RPE-1 cells and CRISPR/Cas9 gene-edited NIH3T3 cells expressing endogenous levels of EHD1-GFP (Fig. 6I and N). As previously reported, depletion of one retromer core complex subunit also decreased expression of the other subunit (Arighi et al., 2004), consistent with the notion that the complex becomes destabilized when any subunit is missing. Significantly, reduced expression of either VPS26 or VPS35 did not alter recruitment of MICAL-L1 to centrioles in RPE-1 cells (Fig. 6F-H; quantified in Fig. 6J) or EHD1-GFP to centrioles in NIH3T3 cells (Fig. 6K-M; quantified in Fig. 6O).

### VPS26 and VPS35 are required for CP110 removal from the mother centriole to facilitate ciliogenesis

A key step in the process of primary ciliogenesis is the removal of the centriolar capping protein, CP110, from the mother centriole, thus facilitating the assembly of ciliary vesicles (Lu et al., 2015). To further our understanding of the mechanism by which the retromer regulates ciliogenesis, we asked whether the retromer plays a role in removal of CP110 from the mother centriole during the early stages of ciliogenesis and depleted VPS35 and VPS26 from RPE-1 cells with siRNA oligonucleotides (Fig. 7E). Indeed, whereas ∼80% of Mock-treated cells displayed CP110 loss from the mother-centriole (Fig. 7A, see dashed oval; quantified in D), CP110 removal was detected in less than 60% of cells depleted of VPS26 (Fig. 7B; quantified in Fig. 7D) or VPS35 (Fig. 7C; quantified in D). Although the decrease in CP110 removal was typically about 25%, it nonetheless suggests a role for the retromer in a relatively early stage of ciliogenesis. Moreover, consistent with our previous findings that neither EHD1 nor MICAL-L1 is required for the recruitment of myosin Va-positive preciliary vesicles to the distal appendages (Xie et al., 2019), depletion of neither VPS26 nor VPS35 had an effect on preciliary vesicle movement to the centriolar distal appendages (Supplemental Fig. 2A-C; quantified in Supplemental Fig. 2D).

**Fig. 7.**
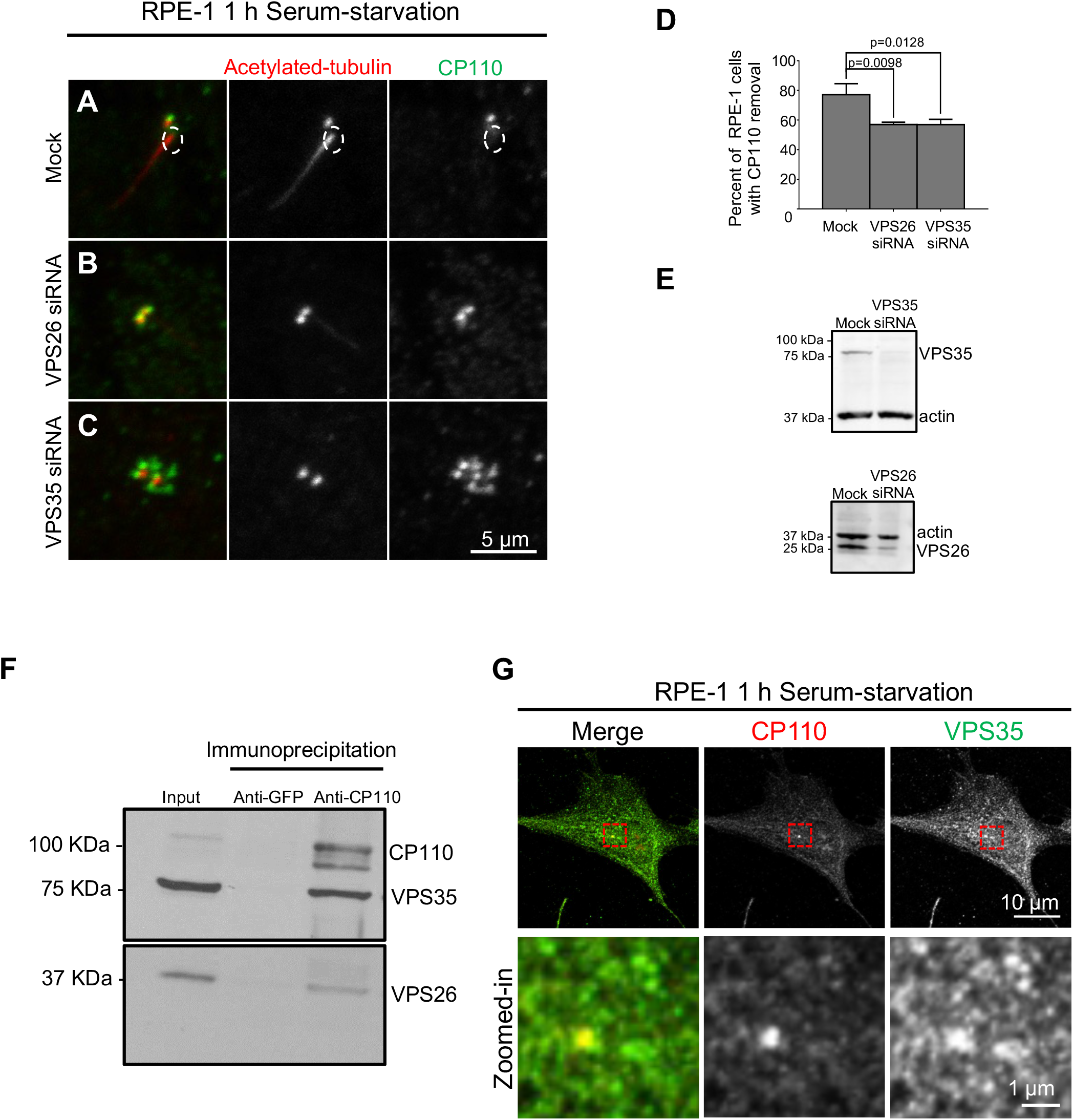
VPS26 and VPS35 regulate CP110 removal from the mother centriole prior to ciliogenesis. (A-E) Mock-treated (A), VPS26-siRNA-treated (B), or VPS35-siRNA-treated (C) RPE-1 cells were serum starved for 1 h, fixed and immunostained with antibodies targeting acetylated-tubulin (red) and CP110 (green). CP110 removal from the mother centriole was impaired upon depletion of VPS26 and VPS35. Bar graph (D) shows the percentage of RPE-1 cells displaying CP110 removal from the m-centriole. *n*=3 experiments (>100 cells quantified for each experiment). Error bars denote standard deviation. VPS35- and VPS26-depletion by siRNA was demonstrated by immunoblotting (E). (F) CP110 interacts with VPS26 and VPS35. RPE-1 cell lysates were subjected to immunoblotting (Input) or immunoprecipitation using anti-CP110 antibodies or rabbit anti-GFP antibodies (negative control). Immunoprecipitated proteins were immunoblotted with anti-CP110, anti-VPS35, and anti-VPS26 antibodies. (G) CP110 displays partial co-localization with VPS35 on the mother centriole. RPE-1 cells were fixed and immunostained using anti-CP110 antibodies (red) and anti-VPS35 antibodies (green).

While the mechanisms by which CP110 is removed from the mother centriole upon ciliogenesis are complex and poorly delineated, we aimed to increase our knowledge of the mode by which the retromer might potentially regulate this step. Accordingly, we hypothesized that the retromer might associate with CP110. To this end we performed immunoprecipitations of cell lysates using antibodies directed to CP110, and we immunoblotted with antibodies against VPS35 and VPS26. Intriguingly, we detected bands representing both VPS35 and VPS26, but these bands were absent when a control anti-GFP antibody was used for immunoprecipitation (Fig. 7F), suggesting that CP110 can reside in a complex with the retromer. In addition we also showed that a pool of endogenous VPS35 localized to CP110-labeled centrioles (Fig.7G), further supporting the notion that the retromer forms a complex with CP110 to facilitate its removal from the mother centriole.

## Discussion

Although initially characterized for its role in retrieval of the yeast Vps10p (Seaman et al., 1997) and the mammalian cation-independent mannose-6-phosphate receptor to the Golgi complex (Arighi et al., 2004), recent studies suggest that retromer core complex subunits play crucial roles in regulating neuronal mitochondria function, and mutations in the VPS35 subunit have been linked to Parkinson’s disease (Braschi et al., 2010; Follett et al., 2014; Vilarino-Guell et al., 2011; Zimprich et al., 2011). Advances in understanding the retromer complex have come from cryo-electron tomography studies and provide evidence for a scaffolding structure in which the VPS26, VPS35 and VPS29 core subunits form an arch that extends from the membrane and the VPS5 subunit (the yeast homolog of mammalian SNX1) assembles on the lipid bilayer of the endosome (Kovtun et al., 2018). However, unlike in yeast, the mammalian retromer may have evolved to function more loosely in coordination with the SNX dimers so that the cation-independent mannose-6-phosphate receptor requires involvement of the SNX1/SNX2-SNX5/SNX6 heterodimeric ESCPE-1 complex (Evans et al., 2020; Simonetti et al., 2019), although this remains somewhat controversial (Seaman, 2021).

While a recent study has elucidated the structure of the metazoan and fungal retromer complexes (Leneva et al., 2021), and various new studies have addressed the retromer in *Drosophila* (Walsh et al., 2021; Ye et al., 2020), fewer studies have addressed retromer function in invertebrate organisms such as *C. elegans*. We chose to address the role of the worm VPS-26 protein based on its high level of amino acid identity with its mammalian VPS26 counterpart (58% identity to the *Homo sapiens* protein). Moreover, since knockdown of a single retromer core complex subunit leads to the degradation of the other subunits (Arighi et al., 2004) and given the homology between retromer core complex structures of different organisms (Leneva et al., 2021), it is fair to assume that knockout of *C. elegans vps-26* gene leads to a dysfunctional worm retromer complex. We found that VPS-26 and the retromer likely regulate worm development, as the number of progeny produced per worm was dramatically reduced by more than 10-fold, potentially as a result of impaired vulva formation often marked by a protruding vulva (Fig. S1). It is noteworthy that the *C. elegans* hermaphrodite vulva is considered a unique model for the study of signaling molecule trafficking within the cell (Schmid and Hajnal, 2015), leading us to speculate that retromer-controlled protein transport might explain the vulval defects seen in *vps-26* knockout worms.

Our studies with a *vps-26* deletion allele generated using CRISPR/Cas9 editing demonstrate that the complete knockout of *vps-26* does not decrease embryonic viability as compared with the control strain (Fig. 3B). However, a previous study which investigated the role of the retromer complex in regulating Wnt gradient formation in *C. elegans* reported in the supplementary data section that the *vps-26(tm1523)* mutant strain exhibits 32% embryonic lethality (Coudreuse et al., 2006). The *tm1523* mutation causes a premature stop codon in the VPS-26 protein, leaving the N-terminal 72 amino acids intact. Thus, this mutation is expected to be a strong loss of function mutation, if not a null mutation. The differences in the embryonic lethality data obtained with the two strains could be explained by potential background mutations that could be present in the *vps-26(tm1523)* strain resulting from the mutagenesis used to generate the knockout allele (Consortium, 2012). Thus, our data highlight the advantages of using CRISPR/Cas9 editing to unambiguously establish *C. elegans* mutant phenotypes.

Another significant developmental difference observed in *vps-26* knockout worms was in body length, with the knockout worms consistently displaying shorter body lengths (Fig. 4). Interestingly, knockdown of the retromer subunit protein VPS-35 in *Xenopus tropicalis* using morpholinos also resulted in decreased body length (Coudreuse et al., 2006), suggesting that the retromer may play an evolutionarily conserved role in maintaining proper body length. Intriguingly, a smaller worm body length has been previously associated with ciliogenesis defects (Fujiwara et al., 2002). Knockout of other related endocytic regulatory proteins, such as SNX3, similarly lead to decreased brood size and worm length (Vieira et al., 2018). Future studies should be directed towards dissecting the mechanism by which the retromer complex functions to maintain a normal body length.

Low brood numbers, such as we observed in our *vps-26* knockout worms, can also be correlated with defects in ciliogenesis, albeit in a 3D culture environment (Lee et al., 2016). Moreover, a recent study indicates that ciliogenesis in *C. elegans* BAG sensory neurons is coordinated by retromer-mediated vesicular trafficking (Martinez-Velazquez and Ringstad, 2018). Two recent studies in mammalian hint at the potential involvement of the retromer complex in ciliogenesis. A study in mammalian cells demonstrated that the retromer complex proteins interact with Polycystin 2, a ciliary protein whose defects are associated with autosomal dominant polycystic kidney disease (Tilley et al., 2018). Another study demonstrated that SNX17, a sorting nexin that binds to the retromer-interacting retriever complex and is involved in sorting, fission and recycling at the endosome (Dhawan et al., 2020; Donoso et al., 2009; Steinberg et al., 2012; Stockinger et al., 2002), also regulates ciliogenesis (Wang et al., 2019). Collectively, these findings suggest that the retromer may regulate ciliogenesis and led us to directly address this question in mammalian cells.

Our data demonstrate a greater than two-fold reduction in ciliated mammalian RPE-1 cells upon serum starvation when either VPS26 or VPS35 are targeted by siRNA oligonucleotides. Coupled with additional data showing that depletion of SNX1 also dramatically reduces primary ciliogenesis, it is evident that both the retromer core complex proteins and the affiliated sorting nexin ESCPE-1 complex contribute to the process. While the VPS26 depletion appears to cause an even more significant reduction in ciliated cells than VPS35 depletion, we anticipate that these differences do not necessarily represent any hierarchy in significance of the retromer proteins, but more likely reflect differential efficiency of retromer core complex subunit depletion by siRNA. Moreover, knocking down one core complex subunit is known to affect expression of the other subunits (Arighi et al., 2004), further complicating any attempts to assign significance to the modest differences observed in ciliogenesis between VPS26 and VPS35 upon their depletion. Overall, however, the data point to a clear requirement for both the retromer core complex and its affiliated SNX dimer/ESCPE-1 complex for normal primary cilia biogenesis, highlighting a common role for both protein assemblages in mammalian cells, at least in the regulation of primary cilia formation.

A number of studies have identified interactions between retromer complex members and other endocytic regulatory proteins, including EHD1 and MICAL-L1 (Gokool et al., 2007; McKenzie et al., 2012; Zhang et al., 2012a; Zhang et al., 2012b), and the latter two proteins have been clearly implicated in ciliogenesis and/or localized to the centrosome (Naslavsky and Caplan, 2020; Xie et al., 2019; Xie et al., 2018). However, despite these interactions, the retromer and EHD1/MICAL-L1 appear to be recruited to the basal body/centrioles independently of one another. MICAL-L1 is recruited to the centrosome through its interactions with microtubules, and in turn recruits EHD1 (Xie et al., 2019). How the retromer is independently recruited remains to be determined, and potential mechanisms may include through the multitude of proteins that interact with the core complex subunits, in particular VPS35.

Although the retromer is recruited to the centrosome/basal body independently of EHD1, the retromer acts at an early stage of ciliogenesis similar to EHD1 and MICAL-L1. Removal of the CP110 capping protein from the mother centriole is a critical step in ciliogenesis, and Rab11, EHD1 and MICAL-L1 are required to facilitate this event (Feng et al., 2012; Feng et al., 2015; Lu et al., 2015; Xie et al., 2019), whereas Rab8 plays a role later on post-CP110 removal (Knodler et al., 2010). While the mechanism for CP110 removal is complex and may be regulated by multiple pathways (Goncalves et al., 2021; Hossain et al., 2017; Kobayashi et al., 2011; Lo et al., 2019; Spektor et al., 2007), the interaction we have observed between the retromer core complex proteins VPS26 and VPS35 with CP110 hints at involvement of a potential vesicular transport pathway. Given the recent involvement of vesicular trafficking in non-endocytic events (Farmer et al., 2017; Naslavsky and Caplan, 2020; Xie et al., 2018), future studies will be aimed at determining whether the retromer and EHD1 might mediate a vesicular transport system that interacts with CP110 and removes it from the mother centriole.

## MATERIALS AND METHODS

### *C. elegans* growth and maintenance

All *C. elegans* strains were grown on MYOB agar plates seeded with OP50 bacteria and maintained at 20°C. The strains used in this study are listed in Supplemental Table S1.

### CRISPR/Cas9 genome editing

CRISPR/Cas9 editing to generate the *vps-26::ha* (IYR010) and *vps-26* knockout (IYR021) strains was performed using assembled ribonucleoprotein complexes as described previously (Iyer et al, 2019; Smith et al, 2020). The sequences of the primers, guide RNAs and repair templates used for making each strain are listed in Supplemental Table S2. The *vps-26::ha* strain was generated first in the wild-type N2 background. The *vps-26::ha* strain then served as the background for generating the *vps-26* knockout strain. All generated strains were confirmed for homozygosity by PCR or by PCR followed by restriction digestion and agarose gel electrophoresis as well as by DNA sequencing.

### Preparation of *C. elegans* lysates for immunoblotting

*C. elegans* lysates for immunoblotting were prepared as follows. Briefly, 100 gravid adults for each genotype were picked into 1 ml of M9 buffer (3 g KH_2_PO_4_, 6 g Na_2_HPO_4_, 5 g NaCl, 1 ml 1 M MgSO_4_ and H_2_O to 1 liter). The worms were washed twice in 1 ml of M9 buffer by centrifugation at 300g for 5 minutes. The worm pellet was then resuspended in 40 µl of 4X SDS-PAGE sample buffer (0.25M Tris HCl pH 6.8, 8% sodium dodecyl sulfate (w/v), 40% glycerol, 20% β-mercaptoethanol and 0.05% (w/v) bromophenol blue in distilled water), heated at 95°C for 10 minutes and stored at -30°C until further use.

### Immunoblotting of *C. elegans* lysates

*C. elegans* lysates were subjected to immunoblotting using the wet transfer method. Briefly, 10 µl of each worm lysate was loaded onto each well of a 10% Bis-Tris gel. The gels were run at a constant current of 60 to 80 mA for approximately 2 h until sufficient band separation was achieved. The proteins were transferred from the gel onto a 0.2 µm nitrocellulose membrane using the wet transfer method (Bio-Rad Laboratories, Hercules, CA). Overnight transfer was performed for 16 h using a constant voltage of 30V. The membranes were blocked using the Intercept blocking buffer (LI-COR Biosciences, Inc., Lincoln, NE), incubated with rabbit anti-HA (1:1000) (Cell Signaling Technology, Inc., Danvers, MA; catalog #3724S) and mouse anti-tubulin (1:200) (Santa Cruz Biotechnology, Inc., Dallas, TX; catalog # sc-32293) antibodies for 2 h and washed thrice with TBST (1X Tris-Buffered Saline, 0.1% Tween 20: 20 mM Tris, 150 mM NaCl, 0.1% Tween 20). The membranes were then incubated with goat anti-mouse (LI-COR Biosciences, Inc. catalog # 926-68070) and donkey anti-rabbit (LI-COR Biosciences, Inc. catalog # 926-32213) IR-dye secondary antibodies for 1 h, washed thrice with TBST and imaged using the LI-COR Odyssey CLx imager (LI-COR Biosciences, Inc.).

### Immunostaining of *C. elegans* embryos and imaging

Immunostaining of *C. elegans* embryos was performed as described previously (O’Connell and Golden, 2014) with a few modifications. Briefly, gravid adults were dissected in 18 µl of M9 buffer to release embryos. The embryos were subjected to freeze cracking by flash-freezing the slides in liquid nitrogen and flicking off the glass coverslip placed on top of the embryos. The embryos were then fixed in 100% methanol prechilled to -30°C and blocked with TBSBT (1X Tris-Buffered Saline with 5% Bovine Serum Albumin and 0.5% Tween 20). The embryos were incubated with rabbit anti-HA (1:1000) (Cell Signaling catalog #3724S) and mouse anti-alpha-tubulin (1:200) (Santa Cruz Biotechnology, Inc. catalog # sc-32293) antibodies for overnight at 4°C. The slides were washed thrice with TBSBT and incubated with goat anti-mouse Alexa Fluor 568 (ThermoFisher Scientific Inc., Catalog #A-11004) and goat anti-rabbit Alexa Fluor 488 (ThermoFisher Scientific Inc., Waltham, MA; catalog #A-11034) secondary antibodies for 1 h at room temperature. The slides were then washed thrice with TBSBT and the embryos were mounted on glass coverslips using the Vectashield mounting medium containing DAPI (Vector Laboratories, Inc., Burlingame, CA; catalog # H-2000-10). The mounted slides were kept in the dark at room temperature overnight and the coverslips were sealed with nail polish 24 h later. The slides were then imaged using a Yokagawa spinning disk mounted on an Olympus confocal microscope with an xyz automated stage equipped with 3 laser lines (405nm, 488nm, 561nm) (Olympus America, Inc., Center Valley, PA).

### *C. elegans* brood count assays and embryonic viability assays

All brood count assays were performed at 20°C. For brood counts, L4 stage *C. elegans* from each genotype were transferred onto 35 mm MYOB plates seeded with OP50 bacteria. Each L4 worm was transferred onto a single MYOB plate and the plate was numbered. The parent worm from each numbered plate was transferred to a new plate every 24 h until the parent stopped producing progeny. All the progeny produced by each of the parent worms were counted each day starting from Day 3 and the number of dead and live progeny were determined. For embryonic viability assays, the percentage of live progeny produced by each parent worm was quantified by dividing the number of live progeny produced by each worm by the total number of progeny produced by the same worm.

### *C. elegans* body length measurements and vulva imaging

Differential Interference Contrast (DIC) live imaging was performed using a Nikon Eclipse fluorescence microscope (Nikon Instruments, Inc., Melville, NY) to measure *C. elegans* body length and to examine *C. elegans* vulva development. For these assays, gravid adult worms of each genotype were picked into 6 to 8 µl of M9 buffer with 1 mM Levamisole. The worms were then mounted on a 2% agarose pad and imaged at either a 20X (body length measurement assays) or a 40X magnification (vulva development assays) using the Nikon NIS-Elements software (Nikon Instruments, Inc.). For *C. elegans* body length measurements, multiple images were taken for each worm at 20X magnification. The images were then aligned, overlapped and stitched together using the Adobe Photoshop software (Adobe Inc., Mountain View, CA). The stitched images were used to measure worm body length using the Nikon NIS-Elements software (Nikon Instruments, Inc.).

### Statistical analysis of *C. elegans* studies

For *C. elegans* brood size, embryonic viability and body length measurement assays, the data obtained were used to generate graphs using the GraphPad Prism software (GraphPad Software, Inc., San Diego, CA). A two-tailed unpaired t-test was used to calculate the p-value for each assay to determine if the observed differences between the two genotypes were statistically significant. Error bars represent the standard deviation.

### Reagents and antibodies for mammalian cell studies

For catalog number and dilutions, please see Table S3. Antibodies were purchased from suppliers as indicated: mouse monoclonal anti-acetylated-tubulin antibodies (Sigma-Aldrich, St. Louis, MO), rabbit polyclonal antibodies to anti-acetylated-tubulin (Cell Signaling, Danvers, MA), rabbit polyclonal anti-CP110 antibodies (ProteinTech, Rosemont, IL), rabbit polyclonal anti-myosin Va antibodies (Novus Biologicals, Littleton, CO), mouse monoclonal anti-GFP antibodies (Sigma-Aldrich, St. Louis), mouse monoclonal anti-MICAL-L1 antibodies (Novus Biologicals), mouse monoclonal anti-glyceraldehyde 3-phosphate dehydrogenase (GAPDH) antibodies conjugated with HRP (ProteinTech), mouse monoclonal anti-actin antibodies (Novus Biologicals), rabbit polyclonal anti-SNX1 antibodies (Novus Biologicals), rabbit polyclonal anti-VPS26 antibodies (Abcam, Cambridge, MA), rabbit monoclonal anti-VPS35 antibodies (Abcam). All secondary antibodies used for immunofluorescence were purchased from Molecular Probes (Eugene, OR).

### Cell culture, induction of ciliogenesis and siRNA knockdown

The human epithelial cell line hTERT RPE-1 (ATCC-CRL4000) was grown at 37°C in 5% CO_2_ in DMEM/F12 (Thermo Fisher Scientific, Carlsbad, CA) containing 10% fetal bovine serum (FBS; Sigma-Aldrich), 2 mM L-glutamine, 100 U/ml penicillin/streptomycin and 1X non-essential amino acids (ThermoFisher Scientific, Waltham, MA). The CRISPR/Cas9 gene-edited NIH3T3 cell line expressing endogenous levels of EHD1 with GFP attached to its C-terminus was generated as described previously (Yeow et al., 2017). Gene-edited NIH3T3 cells were cultured at 37°C in 5% CO_2_ in DMEM (GE Healthcare, Chicago, IL) containing 10% FBS, with 2 mM L-glutamine and 100 U/ml penicillin/streptomycin. To induce ciliogenesis, RPE-1 cells were incubated in DMEM/F12 with 2 mM L-glutamine, 1X non-essential amino acids and 0.2% FBS for the indicated times. NIH3T3 cells were grown in serum-free DMEM with 2 mM L-glutamine for the indicated times.

For siRNA knockdown, ∼5.0×10^4^ RPE-1 or NIH3T3 cells were plated on coverslips in a 35 mm culture dish for 16 h, and then transfected with 200 nM of oligonucleotides using Lipofectamine RNAi/MAX (Invitrogen, Carlsbad, CA) for 72 h in the absence of antibiotics, as per the manufacturer’s protocol. Oligonucleotides targeting human VPS26a (5’-GCCUUACCUGGAGAACUGA[dT][dT]-3’), human VPS35 (5’-GGUGUAAAUGUGGAACGUU[dT][dT]-3’), human EHD1 (5’-CCAAUGUCUUUGGUAAAGA[dT][dT] -3’), human MICAL-L1 (5’-GACAAUGUCUUCGAGAAUA(dT)(dT)-3’), mouse VPS26a (5’-CUAUUCCGAUAAGAUUGUU[dT][dT]-3’&5’-GAAUUACAGCUGAUCAAGA[dT][dT]-3’), and mouse VPS35 (5’-GAUUCGAGAAGAUCUCCCA[dT][dT]-3’&5’-GUAAUGUUCUGGAUUAUAA[dT][dT]-3’) were purchased from Sigma-Aldrich. At 24 h after transfection, the culture medium was replaced with fresh starvation medium for ciliogenesis induction. The efficiency of knockdown was determined by immunoblotting.

### Co-immunoprecipitation

hTERT RPE-1 cells growing in 100 mm dishes were lysed in lysis buffer containing 50 mM Tris-HCl pH 7.4, 150 mM NaCl, 1 mM MgCl_2_, 1% Triton-X and freshly added protease inhibitor cocktail (Roche, Basel, Switzerland). Cell debris was eliminated by centrifugation at 1889 ***g*** at 4°C for 10 min. The cleared lysate was collected and incubated with anti-CP110 antibodies overnight at 4°C. Protein G-agarose beads (GE Healthcare) were added to the lysate–antibody mix and left to rock at 4°C for 2 h. Samples were then washed three times with washing buffer containing 50 mM Tris-HCl pH 7.4, 150 mM NaCl, 1 mM MgCl_2_ and 0.1% Triton X-100. Protein complexes were eluted from the beads by boiling the sample for 10 min in the presence of 4X loading buffer containing 250 mM Tris-HCl pH 6.8, 8% SDS, 40% glycerol, 5% β-mercaptoethanol and 0.2% (w/v) Bromophenol Blue. Eluted proteins were detected by immunoblotting.

### Immunoblotting of proteins from mammalian cells

Cells in culture were washed twice with pre-chilled PBS and harvested with a rubber cell scraper. Cell pellets were lysed by resuspending in lysis buffer containing 50 mM Tris-HCl pH 7.4, 150 mM NaCl, 1% NP-40, 0.5% sodium deoxycholate and freshly added protease inhibitor cocktail (Roche) for 30 min on ice. The cell lysates were then centrifuged at 1889 ***g*** at 4°C for 10 min. The concentration of protein from each sample was measured with Bio-Rad protein assay (Bio-Rad, Hercules, CA), equalized, and eluted by boiling with 4X loading buffer. Proteins from either cell lysates or immunoprecipitations were separated by SDS-PAGE on 10% gels, and transferred onto nitrocellulose membranes (Midsci, St Louis, MO). Membranes were blocked for 30 min at room temperature in PBS containing 0.3% (v/v) Tween-20 (PBST) and 5% dried milk, and then incubated overnight at 4°C with diluted primary antibodies. Protein–antibody complexes were detected with HRP-conjugated goat anti-mouse-IgG (Jackson Research Laboratories, Bar Harbor, ME) or donkey anti-rabbit-IgG (GE Healthcare) secondary antibodies for 1 h at room temperature, followed by enhanced chemiluminescence substrate (ThermoFisher Scientific). Immunoblot images were acquired by iBright Imaging Systems (Invitrogen).

### Immunofluorescence and microscopy imaging of mammalian cells

Cells plated on coverslips were fixed in 100% Methanol at -20°C for 5 min, followed by three rinses with PBS buffer. For immunofluorescence staining, cells were first permeabilized in 0.5% Triton-X plus 0.5% BSA in PBS for 30 min, and then stained with appropriate primary antibodies diluted in PBS buffer containing 0.1% Triton-X and 0.5% BSA for 1 h at room temperature. PBS washes were applied to remove unbound primary antibodies. Cells were then incubated with fluorochrome-conjugated secondary antibodies for 1 h at room temperature and washed three more times in PBS. Coverslips were mounted in Fluoromount G Mounting medium (SouthernBiotech). Imaging was performed with a Zeiss LSM 800 confocal microscope (Carl Zeiss, Oberkochen, Germany) using a Plan-Apochromat 63X/1.4 NA oil objective and appropriate filters, as previously described (Xie et al., 2016). Image acquisition was carried out with Zen software (Carl Zeiss). *Z*-slices were *z*-projected with ImageJ (National Institutes of Health, Bethesda, MD). Images were cropped, adjusted for brightness (whole-image adjustment) with minimal manipulation for better presentation. For quantification, three independent experiments were carried out, and the number of samples collected for quantification is described in the text.

### Statistical analysis of mammalian cell experiments

Data obtained from ImageJ was exported to GraphPad Prism 6 (GraphPad, San Diego, CA). Bar graphs were created representing the mean and standard deviation from data obtained from three independent experiments. Statistical significance was calculated with an unpaired two-tailed *t*-test.

## ACKNOWLEDGMENTS

The authors would like to acknowledge the National Institute of General Medical Sciences (NIGMS) of the National Institutes of Health (NIH) for the following grant support [R01GM123557 and R01GM133915 to S.C., 5P20GM103636 to J.I.], TU Student Research Grants [E.S. and C.D.], TU Faculty Summer Development Fellowship [J.I.], TU Chemistry Summer Undergraduate Research Program [E.S., C.D., D.M., A.D.], Tulsa Undergraduate Research Challenge [E.S., C.D., D.M.], and TU start-up funds [J.I.] The authors declare no competing interests.

---

**Table S1.**

| STRAIN | GENOTYPE | METHOD | RESOURCE |
| --- | --- | --- | --- |
| N2 | WT | N/A | CGC |
| IYR010 | *vps-26(luv10[vps-26::ha]) IV* | CRISPR | This study |
| IYR021 | *vps-26(luv21) IV* | CRISPR | This study |

**Table S2.**
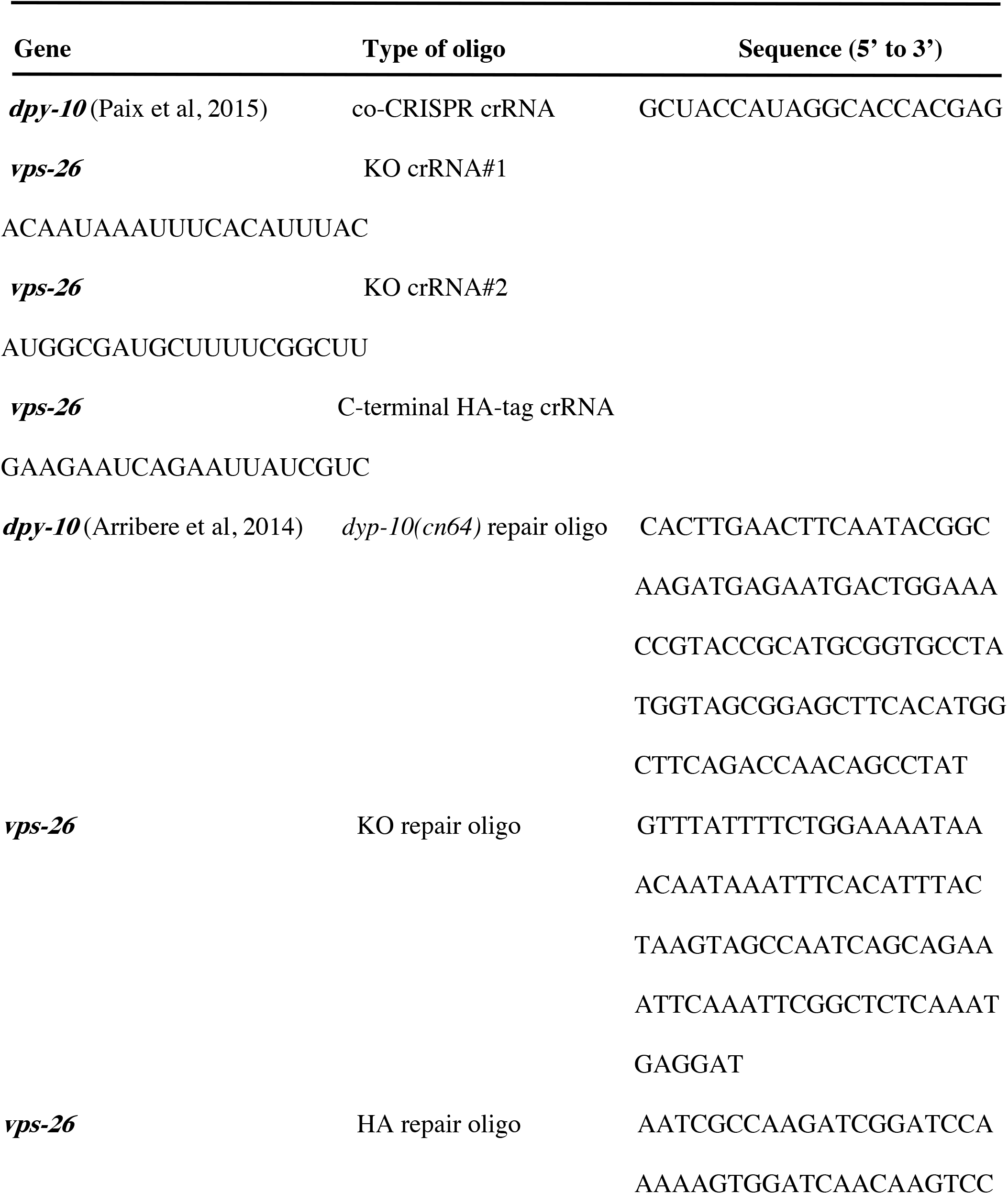
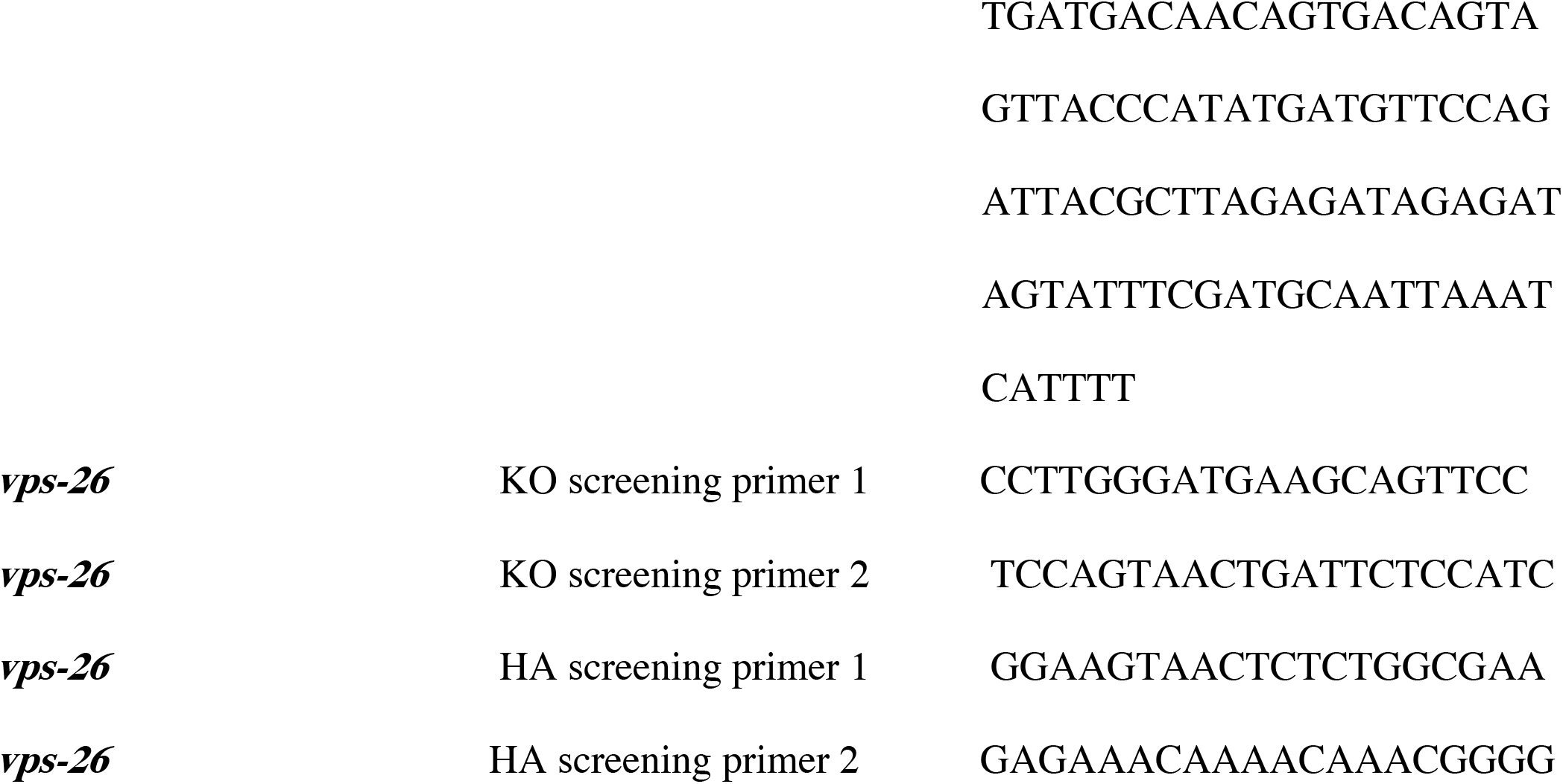

**Table S3.**

| **Antibodies** | **Host** | **Manufacturer** | **Catalogue#** | **Application** | **Dilution** |
| --- | --- | --- | --- | --- | --- |
| **Acetylated-tubulin** | **Mouse** | **Sigma** | **T7451** | **IF** | **1:100** |
| **Acetylated-tubulin** | **Rabbit** | **Cell signaling** | **5335** | **IF** | **1:100** |
| **Alpha tubulin** | **Mouse** | **Santa Cruz Biotechnology** | **sc-32293** | **IB, IF** | **1:200** |
| **CP110** | **Rabbit** | **Protein Tech** | **12794-1-AP** | **IF, IB,IP** | **1:200 (IF)**<br>**1:2000(IB)** |
| **Myosin Va** | **Rabbit** | **Novus** | **NBP1-92156** | **IF** | **1:500** |
| **GFP** | **Mouse** | **Roche** | **1184460001** | **IF** | **1:200** |
| **GFP** | **Rabbit** | **Pierce** | **PA1-980-A** | **IP** |  |
| **HA** | **Rabbit** | **Cell Signaling** | **3724S** | **IB, IF** | **1:1000** |
| **Actin** | **Mouse** | **Novus** | **NB600-535** | **IB** | **1:5000** |
| **MICAL-L1** | **Mouse** | **Novus** | **H00085377-B01P** | **IF** | **1:200** |
| **MICAL-L1** | **Rabbit** | **LifeTein** | **RB1794** | **IB** | **1:1000** |
| **GAPDH-HRP** | **Mouse** | **Protein Tech** | **HRP60004** | **IB** | **1:2000** |
| **VPS26** | **Rabbit** | **Abcam** | **Ab23892** | **IB** | **1:1000** |
| **VPS35** | **Rabbit** | **Abcam** | **Ab157220** | **IF,IB** | **1:200 (IF)**<br>**1:1000 (IB)** |
| **SNX1** | **Rabbit** | **Novus** | **NBP2-13359** | **IB** | **1:500** |
| **Mouse HRP light chain only** | **Goat** | **Jackson** | **115-035-174** | **IB** | **1:7000** |
| **Rabbit HRP** | **Donkey** |  |  | **IB** | **1:5000** |
| **Mouse IRDye 680 RD** | **Goat** | **LI-COR** | **926-68070** | **IB** | **1:14000** |
| **Rabbit IRDye 800CW** | **Donkey** | **LI-COR** | **926-32213** | **IB** | **1:14000** |
| **Mouse Alexa 488** | **Goat** | **Molecular Probe** | **A11029** | **IF** | **1:500** |
| **Rabbit Alexa 568** | **Goat** | **Molecular Probe** | **A11036** | **IF** | **1:500** |
| **Mouse Alexa 568** | **Rabbit** | **Molecular Probe** | **A11061** | **IF** | **1:500** |
| **RabbitAlexa488** | **Goat** | **Molecular Probe** | **A11034** | **IF** | **1:500** |
| **Mouse Alexa 568** | **Goat** | **Thermo Fisher** | **A-11004** | **IF** | **1:1000** |
| **Rabbit Alexa 488** | **Goat** | **Thermo Fisher** | **A-11034** | **IF** | **1:1000** |

